# Gene family expansion and functional diversification of chitinase and chitin synthase genes in Atlantic salmon (*Salmo salar*)

**DOI:** 10.1101/2022.05.05.490710

**Authors:** Matilde Mengkrog Holen, Matthew Peter Kent, Gustav Vaaje-Kolstad, Simen Rød Sandve

## Abstract

**Background:** Chitin is one of the most abundant polysaccharides in nature, forming important structures in insects, crustaceans, and fungal cell walls. Vertebrates on the other hand are generally considered “non-chitinous” organisms, despite having highly conserved chitin metabolism associated genes. Recent work has revealed that the largest group of vertebrates, the teleosts, have the potential to both synthesize and degrade endogenous chitin. Yet little is still known about the genes and proteins responsible for these dynamic processes. Here we used comparative genomics, transcriptomics, and chromatin accessibility data to characterize the repertoire, evolution, and regulation of genes involved in chitin-metabolism in teleosts, with a particular focus on Atlantic salmon.

**Results:** Reconstruction of gene family phylogenies provide evidence for an expansion of teleost and salmonid chitinase and chitin synthase genes after multiple whole-genome duplications. Analyses of multi-tissue gene expression data demonstrated a strong bias of gastrointestinal tract expression for chitin metabolism genes, but with different spatial and temporal tissue specificities. Finally, we integrated transcriptomes from a developmental time series of the gastrointestinal tract with chromatin accessibility data to identify putative transcription factors responsible for regulating chitin-metabolism gene expression (CDX1 and CDX2) as well as tissue-specific divergence in the regulation of gene duplicates (FOXJ2). These transcription factors are also potential regulators of multiple glycosyltransferases being co-expressed with the chitin remodeling genes.

**Conclusion:** The findings presented here add support to the hypothesis that chitin metabolism genes in teleosts play a role in developing and maintaining a chitin-based barrier in the teleost gut and provide a basis for further investigations into the molecular basis of this barrier.

## INTRODUCTION

Chitin is one of the most abundant polysaccharides in nature, serving as the main building block in insect and crustacean exoskeletons as well as forming structural and protective components in fungi. Chitinases and chitin synthases (CHS) are the two major groups of enzymes that have evolved to degrade and synthesize chitin. Decades of work on these enzymes has revealed that bacterial genes encode chitinases that enable bacteria to degrade and utilize chitin as a nutrient source (Cohen-Kupiec and Chet 1998; Beier and Bertilsson 2013), while eukaryotes rich in chitin (i.e. insects, crustaceans and fungi) depend on endogenous chitinases and CHS for normal growth and development (Gooday 1992; Merzendorfer and Zimoch 2003; Zhu *et al*. 2008; Zhang, Zhang, *et al*. 2011; Zhang, L iu, *et al*. 2011; Eichner *et al*. 2015). Curiously, large and highly conserved repertoires of chitinase and CHS genes are also found in vertebrates that do not rely on chitin as a source of nutrition nor possess obvious chitinous body structures such as exoskeletons. Recent experimental work has shown that teleost fish produce chitin in the gastrointestinal tract (GIT) similar to those found in the insect gut epithelium (peritrophic matrix) (Tang *et al*. 2015; Nakashima *et al*. 2018). This realization contradicts the generally held belief that vertebrates are non-chitinous and questions the dogma that chitin does not play an important role in vertebrate physiology.

The function of chitinases in fish has received attention for several reasons. Firstly, chitin is a major component in the natural diets of many fish species, and speculation exists as to whether chitinases could aid in the degradation of chitin to digestible carbohydrates. While fish tissues are known to possess chitinase activity and several fish chitinases have been identified\ (Fänge *et al*. 1976; Lindsay 1984; Gutowska *et al*. 2004; Zhang *et al*. 2012; Koch *et al*. 2014; Teng *et al*. 2014; Gao *et al*. 2017; Ikeda *et al*. 2017), the activity of these enzymes does not seem to correlate with the ability of fish to digest chitin nor with the amount of chitin in their natural diet (Buddington 1980; Lindsay *et al*. 1984; Kono *et al*. 1987; Danulat 1987; Karlsen *et al*. 2017).

Secondly, chitin is present in many fish tissues and structures, such as the developing gut of zebrafish (*Danio rerio*) (Tang *et al*. 2015), the blenny cuticle of *Paralipophrys trigloides* (Wagner *et al*. 1993), the Ampullae of Lorenzini of Chondrichthyes (Phillips *et al.* 2020) and in the scales of parrotfish (*Chlorurus sordidus*), red snapper *(Lutjanus argentimaculatus*), common carp (*Cyprinus carpio*) and Atlantic salmon (*Salmo salar*) (Zaku *et al*. 2011; Tang *et al*. 2015; Rumengan *et al*. 2017), but the role of chitin in these structures is not known. Thirdly, salmonid fish chitinases have been linked to host-parasite interactions during an infestation of salmon louse (*Lepeophtheirus salmonis*), a small crustacean that feeds on the skin, mucus, and blood of salmonids. For example, resistance to salmon lice in Pink salmon (*Oncorhynchus gorbuscha*) has been suggested to be linked to an increased response of host chitinase in larger fish (Sutherland *et al*. 2011), and upregulation of chitinase gene expression together with genes involved in tissue repair and wound healing in lice-infected skin of Atlantic salmon (Robledo *et al*. 2018).

Our understanding of the repertoire and function of chitin degrading enzymes in vertebrates is mostly derived from studies of mammalian genes and proteins. These genes all belong to the glycoside hydrolase 18 family (GH18), which is an ancient multigene family with a conserved *DXXDXDXE* catalytic motif where glutamate represents the catalytic acid (Terwisscha van Scheltinga *et al*. 1996). In mammals, genes encoding these enzymes can be further subdivided into five main groups. Three of these groups have demonstrated enzymatic activity that enables them to break down chitin; chitotriosidase (CHIT1), acidic mammalian chitinase (CHIA), and di-*N*-acetyl-chitobiase (CTBS). CHIT1 and CHIA are hypothesized to have evolved from one common ancestor gene through whole-genome duplication (WGD) in a common vertebrate ancestor (Hussain and Wilson 2013) and can hydrolyze longer chains of chitin into shorter fragments (chitobiose and chitotriose) (Renkema *et al*. 1995; Boot *et al*. 2001). The CTBS group is more distantly related to CHIT1/CHIA and has evolved to hydrolyze shorter, soluble chitooligosaccharides into *N*-acetyl glucosamine monomers (GlcNAc) allowing for complete degradation of chitin. Chitinase-domain containing protein 1 (CHID1) is another chitinase-related group of proteins that is highly conserved in all vertebrates, although the sequence similarity to other GH18 chitinases is low. Human CHID1 (stabilin-1 interacting protein) lacks essential catalytic residues but contains conserved aromatic residues potentially important for saccharide binding (Meng *et al*. 2010). In mammals, but not in all vertebrates, other saccharide-binding chitinases are termed chitinase-like lectins (CHIL). CHIL are non-enzymatic chitinase-like proteins very similar to CHIA and CHIT1, but with active site mutations that render the proteins catalytically incompetent. Human CHIL (OVGP1, CHI3L1, and CHI3L2) are according to phylogenetic analyses of mammalian CHIL predicted to have evolved from gene duplications of ancestral CHIA and HIT1 (Funkhouser and Aronson 2007; Bussink *et al*. 2007). A newly identified group of vertebrate chitinases that does not fit into any of the five mammalian groups is a group called CHIO (Hussain and Wilson 2013). Like CHIL, CHIO is also hypothesized to have evolved from ancestral CHIA and/or CHIT1. Two rounds of whole-genome duplication events specific for teleost (Ts3R) and salmonid fish (Ss4R) have resulted in an amplification of genes that are closely related to this group. There is, however, a lack of systematic effort to characterize the potential for teleost genomes to encode chitin degrading and synthesizing enzymes.

In this paper, we have characterized the evolution and diversification of genes involved in chitin breakdown and synthesis in Atlantic salmon. Using a comparative approach that combines both comparative and functional genomics, we provide an improved understanding of putative protein functions and gene regulation of chitinase and CHS genes in Atlantic salmon. Our results provide a knowledge base for further functional studies of chitin-biology in teleost fish and support the idea that chitin plays a major role in GIT function and physiology.

## MATERIALS AND METHODS

### Phylogenetic analysis

Orthofinder (v.0.3.1) was used to construct orthogroups using the longest protein isoform sequence from gene. Species included in the orthogroups computation were spotted gar (*Lepisosteus oculatus*, LepOcu1), zebrafish (*Danio rerio*, GRCz10), stickleback (*Gasterosteus aculeatus*, BROADS1), Japanese medaka (*Oryzias latipes*, HdrR), pike (*Esox Lucius*, Eluc_V3), rainbow trout (*Oncorhynchus mykiss*, Omyk_1.0), coho salmon (*Oncorhynchus kisutch*, Okis_V1), Atlantic salmon (*Salmo salar*, ICSASG_v2), human (*Homo sapiens*, GRCh38), and house mouse (*Mus musculus*, GRCm38). For each orthogroup, protein sequences were then aligned using MAFFT (v.7) (Katoh and Standley 2013). A maximum likelihood phylogenetic tree was constructed in MEGA7 (Kumar *et al.* 2016) using a neighbor-joining algorithm with a JTT substitution model and 100 bootstrap replicates.

### Tissue expression profiles

See Supplementary Table 1 for more information about species, tissues examined, number of individuals, and where the data are available. Tissue expression profiles from Atlantic salmon (except stomach, pyloric caeca, and midgut), rainbow trout, zebrafish and pike (n = 1 for all tissues except the liver where n = 3 for rainbow trout and n = 4 for zebrafish) were generated from RNA-sequencing (RNA-seq) data as described previously (Lien *et al.* 2016; Gillard *et al.* 2021). In brief, the STAR aligner with default settings (Dobin *et al.* 2013) was used to map RNA-seq reads to the annotated reference genomes and RSEM (Li and Dewey 2011) was used to estimate read counts. Tissue expression data from the stomach, pyloric caeca and midgut of Atlantic salmon (n = 15 for stomach and pyloric caeca, n = 167 for midgut) were generated from previously published RNA-seq data (Gillard *et al.* 2018; Jin *et al.* 2018) following the described method. The RNA-seq data were mapped to the annotated genome (ICSASG_v2) using the STAR aligner, and the read counts were estimated with HTSeq-count (Anders *et al.* 2015). The read counts were transformed to Transcript Per Million Reads (TPM) values normalized for average transcript length and sample size. To get TPM values, the raw gene counts were first divided by the transcript length before dividing by the total library count number. The mean gene expression value was used for the liver, and the median gene expression value was used for the stomach, pyloric caeca, and midgut. The gene expression values were log-transformed (Log_2_(TPM + 1) before further analysis. The RNA-seq data analysis was performed using R (v.3.6.0).

### Gene expression in pyloric caeca of Atlantic salmon spanning the transition from endogenous to exogenous feeding

The expression profiles of chitinases and CHS in pyloric caeca of Atlantic salmon spanning the developmental transition to external feeding were obtained from an RNA-seq dataset available through ArrayExpress under the project number E-MTAB-8306 and was generated as described previously (Jalili *et al*. 2019). Differences in expression levels compared to day 0 was tested by comparing means of expression using a Wilcoxon test with the function “stat_compare_means” in the R-package “Ggpubr” using the default “wilcox.test” parameter. The p-values were adjusted for multiple testing. Genes with low expression (TPM < 1) were removed before the co-expression analysis and quality control of the resulting genes was conducted using the function “goodSamplesGenes” in the “WGCNA” package in R (Langfelder and Horvath 2008) with the argument “verbose = 3”. The co-expression analysis was carried out using the minimum biweight midcorrelation (“bicor”) function in the “WGCNA” package with the argument «maxPOutliers = 0.05» and genes with a correlation value above 0.69 was referred to as co-expressed genes. Gene enrichment of the co-expressed genes was done using KEGG enrichment with the function “kegga” from the “limma” package in R (Ritchie *et al.* 2015) and the argument «species.KEGG = “sasa”» and the universe specified to be only expressed genes. The p-values returned by “kegga” were not adjusted for multiple testing.

### Chromatin accessibility in pyloric caeca

ATAC-seq reads from pyloric caeca of Atlantic salmon were downloaded from ArrayExpress (E-MTAB-9001). Read mapping and ATAC-peak calling was done using BWA (v.0.7.17) (Li and Durbin 2009) and Genrich v.06 (https://github.com/jsh58/Genrich) as described in detail in Bertolotti *et al.* (2020).

### Transcription factor motif enrichment

DNA sequences from open chromatin (i.e. within ATAC-seq peaks) around TSS (1000bp upstream to 200bp downstream) of chitinase- and CHS genes were used for transcription factor motif scan and enrichment. The scan and enrichment was carried out using SalMotifDB (Mulugeta *et al*. 2019), a tool for analyzing putative transcription factor binding sites in Atlantic salmon. Consensus motifs were obtained using the “ggseqlogo” package in R.

## RESULTS

### Phylogenetic analysis of chitinase protein sequences

The annotated Atlantic salmon genome (ICSASG_v2) includes 12 genes with strong homology to mammalian chitinase genes (see Supplementary Table 2 for gene IDs, proteins accession numbers, and names given in this paper) belonging to the family 18 of the glycoside hydrolases, as classified by the carbohydrate-active enzyme (CAZy) database (Drula *et al.* 2022). To investigate the evolutionary history of the chitinase gene family in fish we reconstructed phylogenetic trees of genes within the glycoside hydrolase family 18 orthogroup. The species selection was designed to include vertebrates that have experienced different numbers of whole-genome duplications. All species share the two whole-genome duplications occurring in the ancestor of all vertebrates, and except gar, all fish species share an additional whole genome duplication at the base of the teleost lineage (Ts3R), while Atlantic salmon, rainbow trout and Coho salmon share an additional fourth salmonid-specific whole-genome duplication event (Ss4R) (Figure 1A).

**Figure 1.**
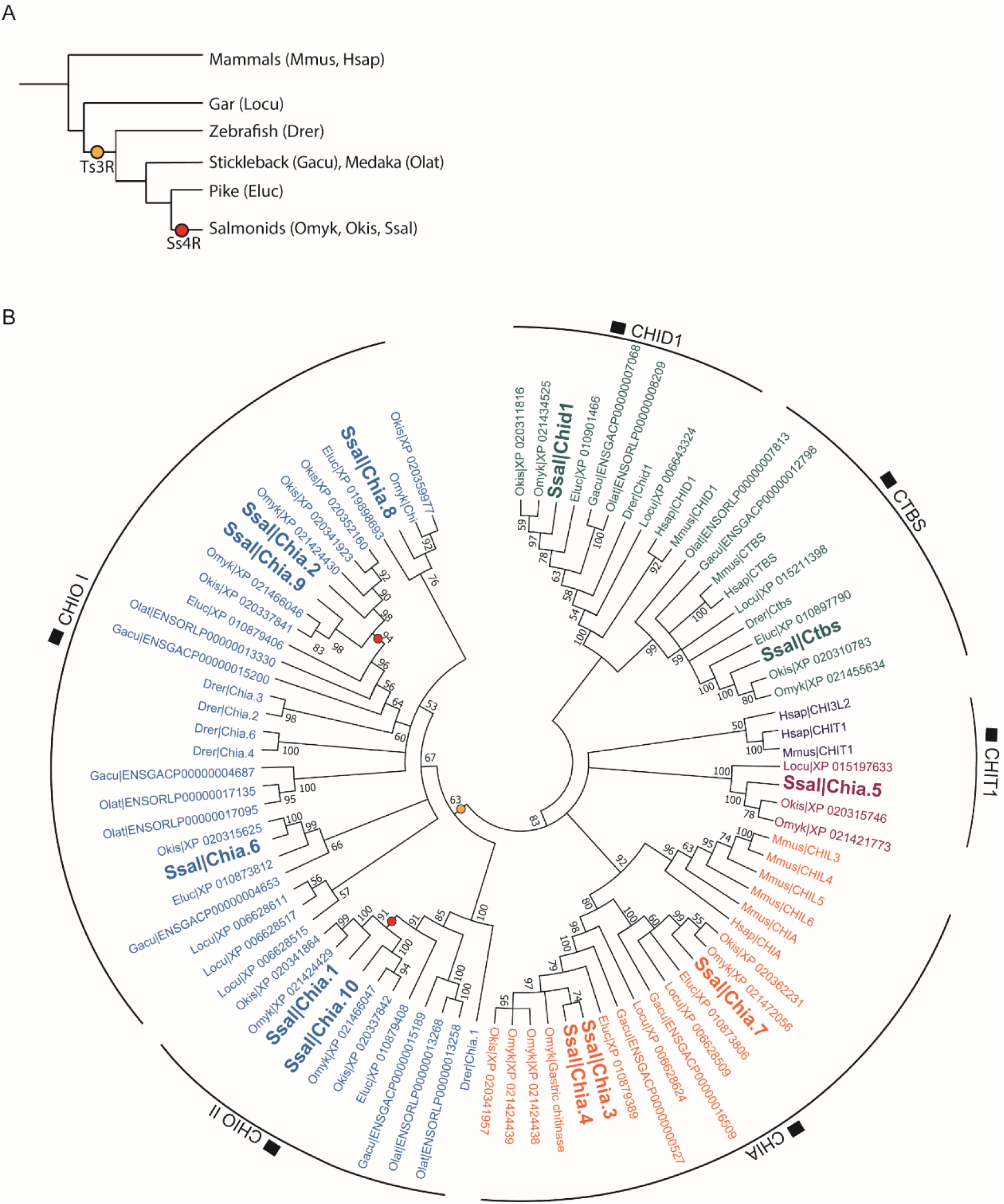
Comparative analysis of GH18 proteins. A) Whole-genome duplication (WGD) events experienced by species included in the phylogenetic comparison. B) Phylogenetic tree of GH18 chitinase proteins in spotted gar (locu), zebrafish (drer), stickleback (gacu), Japanese medaka (olat), pike (eluc), rainbow trout (omyk), coho salmon (okis), Atlantic salmon (ssal), human (hsap) and house mouse (mmus). The colors represent different monophyletic clades in the phylogenetic tree.

Our analysis revealed that the 12 Atlantic salmon chitinase proteins are distributed among six major clades (Figure 1B). These six clades formed two major “superclades” (supported by a high bootstrap value of 83), one containing CTBS and CHID1 type proteins (two from Atlantic salmon) and the other containing CHIT1, CHIA, and CHIO (10 from Atlantic salmon). Of the 10 Atlantic salmon proteins annotated as acidic mammalian chitinases (CHIA) in the NCBI RefSeq annotation (release 100) all salmon proteins share the following chitinase characteristics: a signal peptide, a glycoside hydrolase 18 family catalytic domain, and a chitin-binding domain (CBM14; identified by dbCAN2 annotation (Zhang *et al.* 2018)) at the carboxyl-terminus. Three salmon CHIA proteins (namely Chia.3, Chia.4, and Chia.7) fell into a monophyletic clade (descending from a common ancestor) containing the human acidic mammalian protein AMCase, whereas the remaining seven CHIA protein sequences were distributed among two teleost-specific monophyletic clades. One salmon CHIA protein (Chia.5) will hereafter be referred to as a CHIT1-member. This protein is the only salmon chitinase protein with loss-of-function mutations in the catalytic motif and a truncated chitin-binding domain. The remaining six salmon chitinase proteins (Chia.1, Chia.2, Chia.6, Chia.8, Chia.9, and Chia.10) belong to two clades (termed CHIO I and II) forming a larger monophyletic group.

To make interferences about how ancient whole-genome duplication and other duplication events have contributed to the present diversity of chitinase proteins in teleost fish, the protein sequence phylogeny was compared with the species tree topology. The CHID1 and CTBS clades only contain one protein sequence per species, and the protein trees resemble to a large extent the species topology except for the polytomy in the CTBS clade that fails to place the mammals as a sister clade to the teleost species. This is in agreement with the hypothesis that CTBS and CHID1 genes resulted from an ancient gene duplication before the vertebrate diversification (Funkhouser and Aronson 2007; Hussain and Wilson 2013), possibly the whole genome duplication at the base of all vertebrates. The two distinct fish-specific CHIA subclades are more closely related to each other than to their sister subclade containing the mammalian CHIA proteins. Furthermore, since both fish subclades contain a predicted gar protein it is likely that these fish-specific duplicates arose through a duplication event prior to the divergence of teleosts. In the fish-specific CHIO clade, comprising 39 protein sequences, the three gar-specific proteins cluster closely together and, because of low bootstrap values (< 70) for key splits in the tree, we cannot firmly place these in relation to the remaining teleosts. However, based on the sequence relationships between the teleost CHIO species it is likely that Ts3R has contributed to at least one CHIO-duplication event as previously hypothesized (Hussain and Wilson 2013). Two nodes reflecting the Ss4R event can be inferred in the CHIO I and II clades, including the branches containing Chia.1+10, and Chia.2+9. These proteins are located on homologous regions of different chromosomes (22 and 12) in Atlantic salmon (Supplementary Figure 1).

### Tissue-specific regulatory divergence of chitinase genes in the gastrointestinal tract

A comparative analysis of tissue expression in zebrafish, pike, rainbow trout, and Atlantic salmon (Figure 2) was performed to characterize the divergence of gene regulation encoding chitinase enzymes. Members of the CHIT1 group were not included in this analysis as their expression levels were low, indicating that they may represent pseudogenes encoding non-functional enzymes. Across all species, the results showed a clear bias towards gene expression in GIT and revealed both conserved expression divergence among orthologs in different species as well as lineage-specific regulatory divergence.

**Figure 2.**
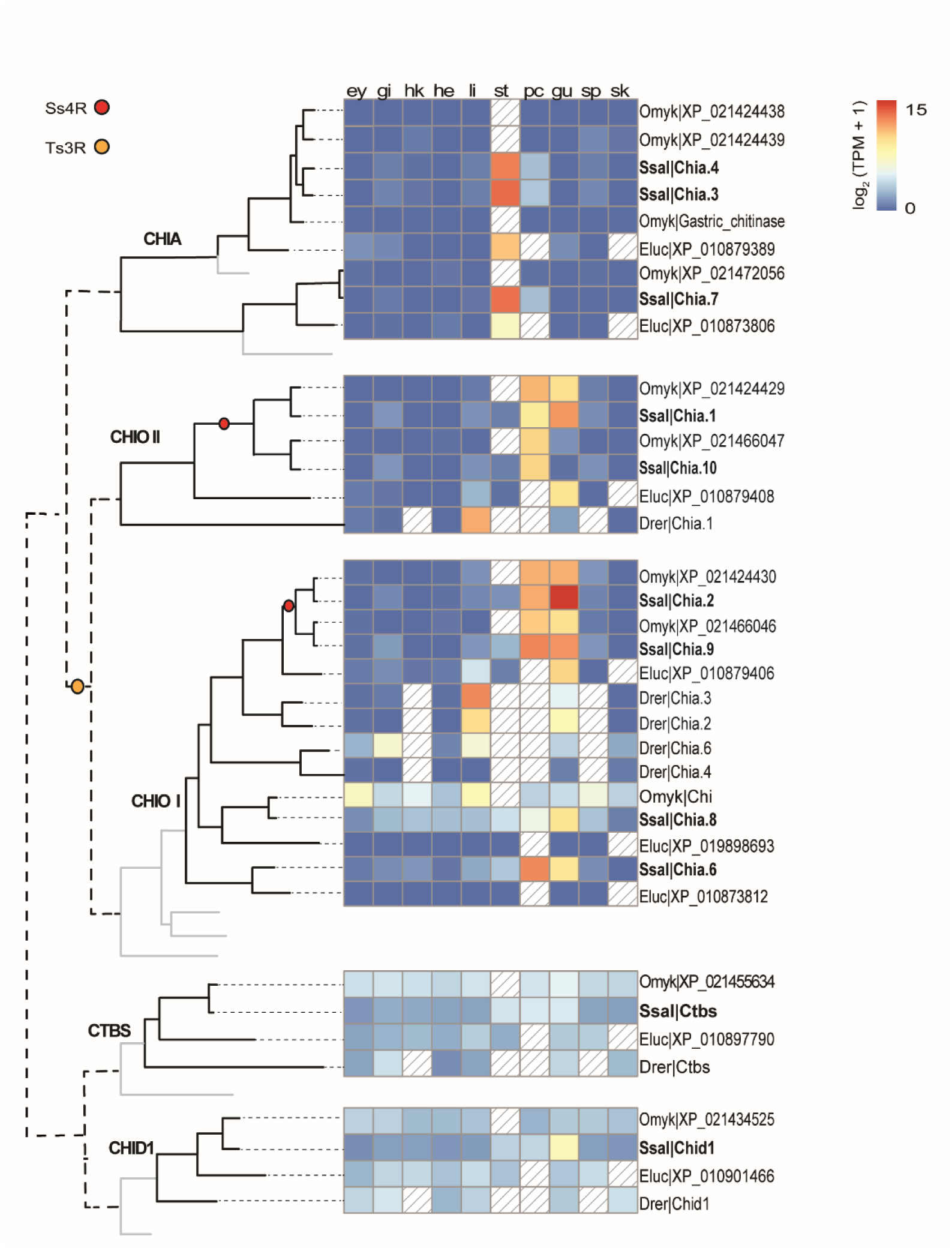
Comparative tissue expression of GH18 genes in zebrafish (drer), pike (eluc), rainbow trout (omyk), and Atlantic salmon (ssal). n≥1. The phylogenetic tree is a simplified version of figure 1B and the lines do not represent real evolutionary distances. Yellow and red circles represent the teleost-specific whole-genome duplication (Ts3R) and salmonid-specific whole-genome duplication (Ss4R), respectively. Solid light grey lines indicate the phylogenetic position of a spotted gar, but expression data were not analyzed for this outgroup. The tissue expression panel shows gene expression of GH18 genes in the following tissues: ey = eye, gi = gill, hk = head kidney, he = heart, li = liver, st = stomach, pc = pyloric caeca, gu = midgut, sp = spleen and sk = skin. Colored boxes indicate gene expression in the range of 0 to 15 log_2_(TPM + 1) values, while diagonal lines represent missing data.

CHIA genes displayed the most conserved tissue expression regulation across all species with stomach-specific expression. A similar stomach bias is also observed for CHIA in mice, bats, pigs, chickens, and humans indicating that CHIA enzymes share an important gastric function that is conserved across fish, mammals, and birds (Boot *et al*. 2001; Strobel *et al*. 2013; Ohno *et al*. 2016; Tabata, Kashimura, Wakita, Ohno, Sakaguchi, Sugahara, Kino, *et al*. 2017; Tabata, Kashimura, Wakita, Ohno, Sakaguchi, Sugahara, Imamura, *et al*. 2017; Tabata *et al*. 2019). Notably, the agastric (stomach-less) zebrafish do not express genes related to gastric functions, including CHIA genes.

The tissue expression profiles of CHIO-I and II genes show different patterns compared to the CHIA genes. Although CHIO genes also display GIT expression dominance, these genes are not stomach-specific but rather expressed in other GIT sections such as pyloric caeca and midgut. Additionally, CHIO gene expression is generally less tissue-specific and has larger inter- and intra-tissue specific variations in tissue expression patterns. For example, while both salmon CHIO II genes are expressed almost exclusively in the GIT, the CHIO I clade contains salmon genes expressed in the GIT and a gene (*chia*.*8*) that is lowly expressed in all tissues examined. We also observe some less striking, but clear, cases of regulatory divergence of CHIO II genes following the more recent Ss4R, with one duplicate being mostly expressed in pyloric caeca (*chia*.*10*), while the other is expressed in both pyloric caeca and gut (*chia*.*1*).

Finally, the CTBS and CHID1 gene groups show a different tissue regulation pattern than the other chitinases. Both gene groups are generally expressed at low levels compared to their CHIA and CHIO counterparts. CHID1 orthologs in zebrafish, pike, and rainbow trout are ubiquitously expressed in all tissues, but the salmon *chid1* gene is expressed at two-fold higher levels in the midgut compared to the stomach and pyloric caeca, and four-fold higher in the midgut compared to non-GIT tissues indicating some tendency to GIT specific expression. Similar to CHID1, fish CTBS are expressed across all tissues, but with two-fold higher expression in the GIT of Atlantic salmon.

### Chitin synthase genes are mainly expressed in pyloric caeca and midgut of teleost fish

The phylogenetic analysis of the gene family containing genes encoding CHS proteins showed a split into two major subclades (I and II) which, since gar has a single gene copy, likely arose in the Ts3R whole-genome duplication event (Figure 3). Furthermore, the salmonid-specific gene copies in subclade II (i.e. *chs1a* and *chs1b* from Atlantic salmon) likely originate from the Ss4R as they are located on chromosomes (28 and 1 respectively) matching the well described synteny within the duplicated Atlantic salmon genome (Lien *et al.* 2016) (Supplementary Figure 1). The tissue expression pattern shows that the CHS genes in subclade I are expressed in low abundance in all tissues, whereas CHS genes in subclade II follow the same expression pattern as CHIO-I and II genes (Figure 2), with expression specific to pyloric caeca and gut (Figure 3).

**Figure 3.**
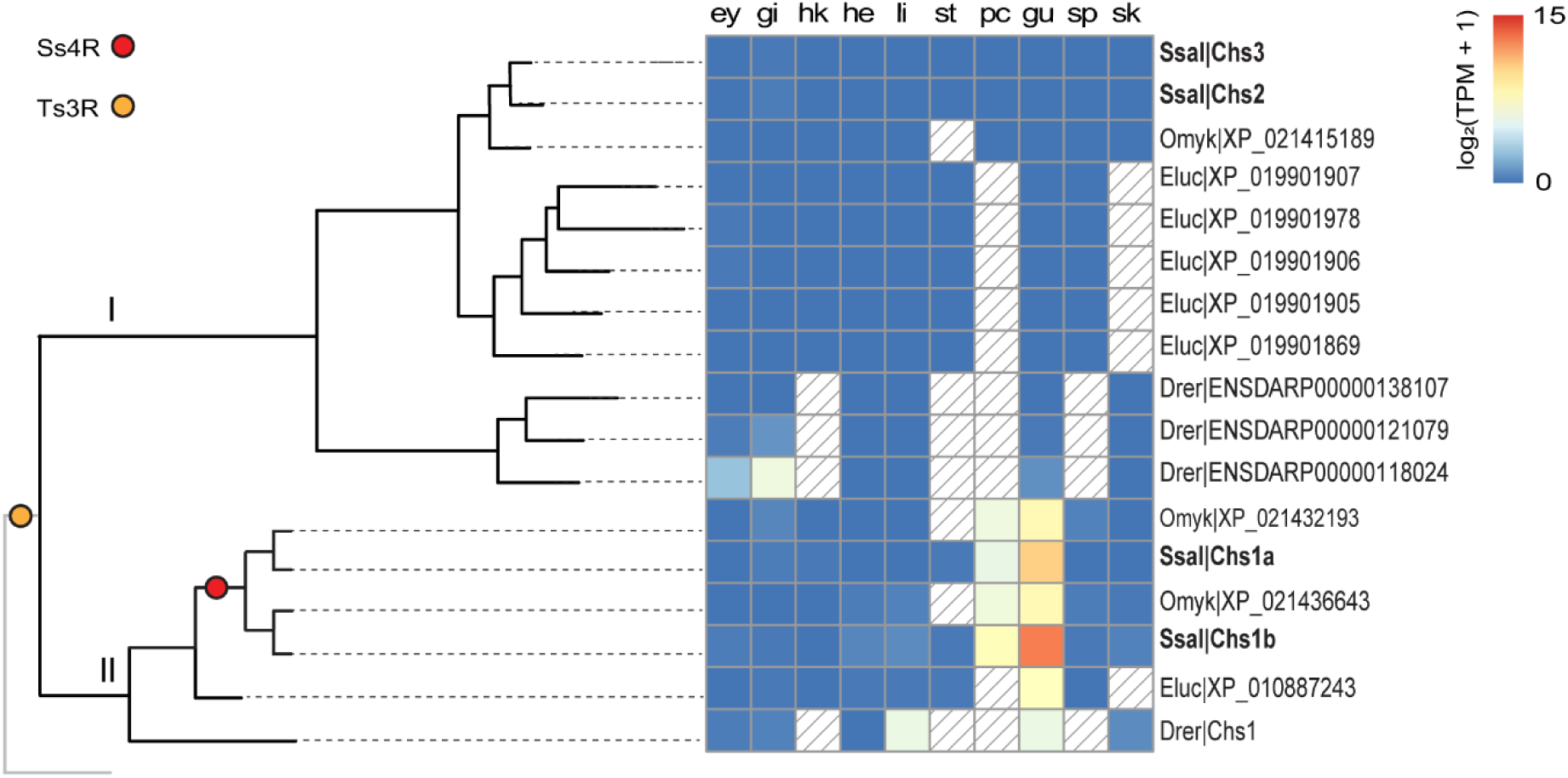
Comparative tissue expression of CHS (chitin synthase) genes in zebrafish (drer), pike (eluc), rainbow trout (omyk), and Atlantic salmon (ssal). n≥1. Yellow and red circles in the illustrative phylogenetic tree represent the teleost-specific whole-genome duplication (Ts3R) and salmonid-specific whole-genome duplication (Ss4R), respectively. The lines do not represent real evolutionary distances. Solid light grey lines indicate the phylogenetic position of a spotted gar, but expression data were not analyzed for this outgroup. The tissue expression panel shows gene expression of CHS genes in the following tissues: ey = eye, gi = gill, hk = head kidney, he = heart, li = liver, st = stomach, pc = pyloric caeca, gu = midgut, sp = spleen and sk = skin. Colored boxes indicate gene expression in the range of 0 to 15 log_2_(TPM + 1) values. Boxes with diagonal lines represent missing data.

Notably, predicted protein sequences of teleost CHS genes contain similar conserved amino acid sequence motifs as found in insect CHS proteins. The motifs *EDR* and *QRRRW* are common for all CHS, while *CATMWHXT* and *QKFEY* are signatures of insect CHS (Merzendorfer and Zimoch 2003). *EDR, QRRW*, and *QKFEY* are motifs found in all predicted fish CHS protein sequences examined, but the *CATMWHXT* motif is present in the CHS subclade II only.

### Gene regulation of chitinases and chitin synthases in pyloric caeca of Atlantic salmon

To be able to better understand the regulation of chitin metabolism genes we leveraged a RNA-seq dataset from pyloric caeca of Atlantic salmon (ArrayExpress, E-MTAB-8306) that spans the developmental transition from endogenous to exogenous nutrition (Jalili *et al*. 2019). The changes in gene expression observed across the developmental time series show two major trends; 5 genes significantly increased expression following external feed intake (p < 0.01), while 3 genes did not (Figure 4A).

**Figure 4.**
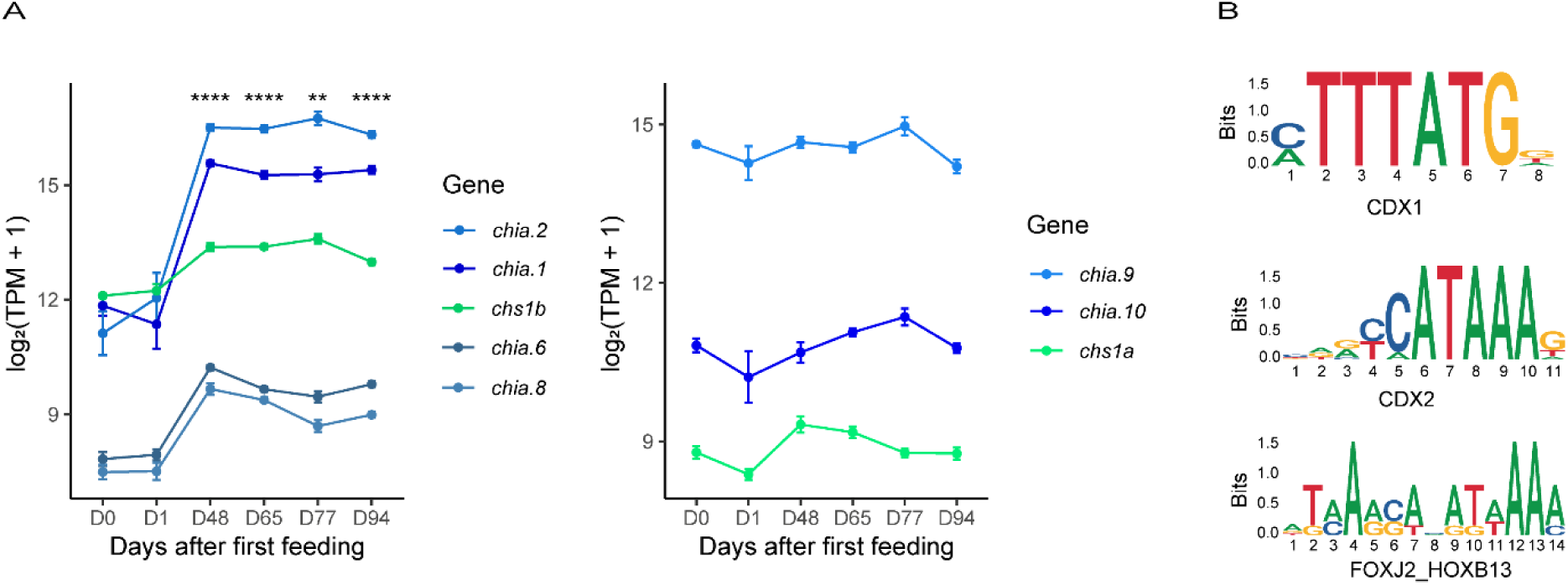
A) Gene expression levels of *chia.1, chia.2, chia.6, chia.8, chia.9, chia.10, chs1a* and *chs1b* before and after external feeding. Chitinase and CHS gene expression in the pyloric caeca of Atlantic salmon days before (D0) and days after external feeding (D1-D94). Please note that the y-axis does not extend to 0. The asterisks (**, ****) indicate a significant difference in expression compared to D0 (p.adj<0.01 and p.adj<0.0001, respectively, n≥4. B) Consensus motifs for binding of transcription factors in promoters of chitinase- and CHS genes. CDX1 and CDX2 motifs are found in promoters of all chitinase- and CHS genes. The FOXJ2_HOXB13 motif is only present in promoters of genes with a significant increase in expression upon transition to external feeding. The numbers indicate the consensus site position of each base, and the vertical axes (Bits) indicate the information content of the base frequency at the given base position.

Co-expressed genes (genes with correlated expression profiles) are often controlled by the same regulators and involved in the same biological processes. To better understand the mechanisms underlying the regulation of genes involved in chitin metabolism, and particularly drive increased expression of some chitinases and CHS following feed ingestion we used a co-expression approach. We first used biweight midcorrelation (bicor) to identify genes with similar expression patterns in the pyloric caeca across the developmental time series. Based on KEGG (Kyoto Encyclopedia of Genes and Genomes) gene enrichment analysis, co-expressed (bicor > 0.69, n = 36) with salmon *chia*.*1, chia*.*2, chia*.*6, chia*.*8*, and *chs1b* were genes involved in metabolic processes like amino sugar and nucleotide sugar metabolism (p-value = 8.79 · 10^−10^) and glycosphingolipid biosynthesis (p-value = 5.92 · 10^−4^) (Supplementary Table 3). We found the chitinase and CHS genes to be associated with the amino sugar and nucleotide sugar metabolism KEGG pathway, together with a UDP-*N*-acetylhexosamine pyrophosphorylase-like gene (*uap1*) which most likely codes for an enzyme that converts uridine triphosphate (UTP) and N-acetylglucosamine-1-phosphate (GlcNAc-1-P) into uridine diphosphate *N*-acetylglucosamine (UDP-GlcNAc) (Mio *et al*. 1998). This is known to result in an activated substrate required for chitin synthesis by CHS. UDP-GlcNAc can also be transferred by beta-1,3-galactosyl-O-glycosyl-glycoprotein beta-1,6-*N*-acetylglucosaminyltransferase (GCNT1) to form mucin-type O-glycan structures. *Gcnt1* is one of the multiple glycosyltransferases being co-expressed with chitinase and CHS genes. Associated with the glycosphingolipid biosynthetic pathway we found two additional glycosyltransferase genes coding for alpha-2,8-sialyltransferase-like proteins, involved in the transfer of sialic acid to produce sialyl glycoconjugates.

To further dissect out putative transcription factors involved in the regulation of chitinase and CHS genes, we performed a transcription factor binding (TFBS) scan for two classes of genes: (1) all CHIO- and CHS genes being highly expressed in pyloric caeca, and (2) those that we find induced during the developmental transition to external feeding exclusively (Figure 4A). We based the scan on open-chromatin regions of promoter sequences of chitinase– and CHS genes. The data show that all chitinase- and CHS genes had open chromatin regions spanning the TSS in pyloric caeca except CHIA-genes and *chia*.*8*. Furthermore, the transcription factor motif scan revealed that two motifs were common for all CHIO- and CHS genes. These motifs were two homeodomain (HOX) related motifs: a caudal type homeobox 1 (CDX1) motif and a caudal type homeobox 2 (CDX2) motif (Figure 4B). For the CHIO-, CHS- and co-expressed genes induced during the transition to external feeding the FOXJ2_HOXB13 motif associated with binding of the forkhead box (FOX) transcriptional factor family (Figure 4B) were enriched (p-value < 0.01).

## DISCUSSION

Our results suggest that Ts3R and Ss4R duplication events resulted in the expansion of chitinase- and CHS genes in fish (Figure 1) and that these genes generally encode proteins with conserved residues in active motifs. This is strikingly different from mammals, which have lost their genes for CHS and where mutations in the active site of chitinases followed by mammal specific gene duplications have resulted in the expansion of non-enzymatic chitinase-like lectins. In general, teleost and salmonid chitinase- and CHS genes share a clear expression bias towards the gastrointestinal tract (Figure 2 & 3). This expression bias may explain the presence of chitin in the gut of zebrafish (Tang *et al*. 2015) and rainbow trout (Nakashima *et al*. 2018), and adds support to a chitin-based mucosal barrier in the gut of teleost fish previously hypothesized to have evolved from the chitin-based barrier we find in invertebrates (Nakashima *et al*. 2018). This is also in line with the increased expression of chitin metabolism genes during development (Figure 4), which coincides with the development of a pyloric caeca with increased complex intestinal mucosal structures (Sahlmann *et al*. 2015). The transition from endogenous to external feeding involves exposure to both larger food particles and new microbial communities. We predict that this exposure boosts the expression of chitinases and CHS needed to synthesize and remodel a chitinous layer that surrounds the intestinal mucosa and protects the intestinal epithelium, in addition to other genes related to intestinal differentiation and mucus production. Co-expression of *uap1* and glycosyltransferases like *gcnt1* supports this assumption, as UAP1 can produce the activated UDP-GlcNAc used by both CHS to produce chitin and by GCNT1. GCNT1 is important for the production of mucins and mice deficient in related genes have been shown to have increased permeability of the mucosal barrier which can alter the mucosal immune homeostasis (Stone *et al*. 2009). Furthermore, the co-expressed alpha-2,8-sialyltransferase orthologs are, in humans, linked to the production of gangliosides; glycosphingolipids that contain one or multiple residues of sialic acid and that is localized in the brush border membrane of intestinal enterocytes. Such gangliosides are known to be important for maintaining intestinal integrity and reducing inflammation (Miklavcic *et al*. 2012).

Little is known about the regulatory networks of chitinases and chitin synthetases. The presence of FOX, CDX1, and CDX2 motifs in the promoter regions of CHIO- and CHS genes presented here support the hypothesis of a possible role in the formation of a chitin-based mucosal barrier. Both CDX1 and CDX2 are known to be major regulators of intestine-specific genes and are crucial for intestinal differentiation. In zebrafish, for example, knockdown and overexpression experiments show that CDX1B, homologous to mammalian CDX1, is responsible for terminal differentiation of goblet cells, cells that are responsible for the secretion of mucins into the intestinal mucosa (Chen *et al*. 2009). Moreover, it is plausible that the FOXJ2_HOXB13 motif could be bound by intestinal FOX proteins like FOXA1 and FOXA2. These proteins are also linked to intestinal goblet cell mucus production (van der Sluis *et al*. 2008). Synergetic transcription factor binding of CDX and FOXA transcription factors has previously been shown to regulate intestine-specific mucins (Jonckheere *et al*. 2007). Similar synergetic effects can possibly explain the difference in gene expression of the chitinase- and CHS genes lacking the FOX motif in their promoter. That said, the chromatin accessibility is likely to change during development and it is important to take into consideration that the ATAC-seq data used to guide the search for transcription factor binding sites in this study was derived from adult fish. Nevertheless, CDX1, CDX2 and FOX are interesting candidates for futures studies into the regulation of chitin metabolism genes in fish GIT.

Functional diversification is observed for the different fish chitinases. We find that, unlike the CHIO- and CHS genes, CHIA genes are exclusively expressed in the stomach and have no open chromatin regions in their promoters in the pyloric caeca of Atlantic salmon. Thus, these genes are likely to be regulated by other transcription factors than the CHIO- and CHS genes. The high degree of sequence similarity to mammalian stomach chitinases suggests that the fish-specific CHIA proteins share an ancestral function related to the presence of an acidic stomach (Krogdahl *et al*. 2015). This hypothesis is strengthened by the loss of CHIA genes in agastric zebrafish. As various fish stomach CHIA proteins have shown to be able to break down chitin structures typically found in the natural diet of Atlantic salmon, like shrimp, squid, and insects (Ikeda *et al*. 2017), we cannot rule out a possible role of teleost CHIA proteins in digestion. The results presented here thus imply that some fish species like Atlantic salmon do have the genetic toolbox needed to tolerate and digest chitin-containing feed.

## CONCLUSION

There has been an expansion of chitinase and chitin-synthase like proteins in Atlantic salmon and different groups of chitinases have evolved to be expressed in different tissues. Based on our results we hypothesize two different roles of Atlantic salmon chitinases and CHS: (1) that stomach-related CHIA proteins aid in the digestion of chitin, whereas (2) CHIO- and CHS proteins are involved in remodelling of chitinous structures surrounding mucosal membranes of pyloric caeca and gut. To verify this, functional characterization of chitinous structures and the enzymes that remodel these are needed. However, this work provides a basis for future functional studies identifying the underlying mechanisms for the presence of chitinous structures in Atlantic salmon.

## DATA AVAILABILITY STATEMENT

The authors declare that the data supporting the results in this study are accessible in the paper and the supplementary file.

## ACKNOWLEDGEMENTS

We want to acknowledge Dr. Gareth Benjamin Gillard, Dr. Yang Jin, and Dr. Thomas Nelson Harvey for their help with bioinformatics.

