## Supplementary Data for "Gene family expansion and functional diversification of chitinase and chitin synthase genes in Atlantic salmon (*Salmo salar*)"

**Supplementary Table 1.** Information about RNA-seq data used to compare tissue expression in different tissues across species.

| **Tissue** | **Species** | **Individuals** | **Project number** | **Database** |
| --- | --- | --- | --- | --- |
| Eye | Zebrafish | 1 | E-MTAB-8959 | ArrayExpress |
| Gill | Zebrafish | 1 | E-MTAB-8959 | ArrayExpress |
| Heart | Zebrafish | 1 | E-MTAB-8959 | ArrayExpress |
| Liver | Zebrafish | 4 | E-MTAB-8959 | ArrayExpress |
| Midgut | Zebrafish | 1 | E-MTAB-8959 | ArrayExpress |
| Skin | Zebrafish | 1 | E-MTAB-8959 | ArrayExpress |
| Eye | Northern pike | 1 | PRJNA221548 | BioProject |
| Gill | Northern pike | 1 | PRJNA221548 | BioProject |
| Head kidney | Northern pike | 1 | PRJNA221548 | BioProject |
| Heart | Northern pike | 1 | PRJNA221548 | BioProject |
| Liver | Northern pike | 1 | PRJNA221548 | BioProject |
| Stomach | Northern pike | 1 | PRJNA221548 | BioProject |
| Midgut | Northern pike | 1 | PRJNA221548 | BioProject |
| Spleen | Northern pike | 1 | PRJNA221548 | BioProject |
| Eye | Rainbow Trout | 1 | E-MTAB-8959 | ArrayExpress |
| Gill | Rainbow Trout | 1 | E-MTAB-8959 | ArrayExpress |
| Head kidney | Rainbow Trout | 1 | E-MTAB-8959 | ArrayExpress |
| Heart | Rainbow Trout | 1 | E-MTAB-8959 | ArrayExpress |
| Liver | Rainbow Trout | 3 | E-MTAB-8959 | ArrayExpress |
| Pyloric caeca | Rainbow Trout | 1 | E-MTAB-8959 | ArrayExpress |
| Midgut | Rainbow Trout | 1 | E-MTAB-8959 | ArrayExpress |
| Spleen | Rainbow Trout | 1 | E-MTAB-8959 | ArrayExpress |
| Skin | Rainbow Trout | 1 | E-MTAB-8959 | ArrayExpress |
| Eye | Atlantic salmon | 1 | PRJNA72713 | BioProject |
| Gill | Atlantic salmon | 1 | PRJNA72713 | BioProject |
| Head kidney | Atlantic salmon | 1 | PRJNA72713 | BioProject |
| Heart | Atlantic salmon | 1 | PRJNA72713 | BioProject |
| Liver | Atlantic salmon | 1 | PRJNA72713 | BioProject |
| Stomach | Atlantic salmon | 15 | PRJEB21981 | European Nucleotide Archive |
| Pyloric caeca | Atlantic salmon | 15 | PRJEB21981 | European Nucleotide Archive |
| Midgut | Atlantic salmon | 167 | PRJEB24480 | European Nucleotide Archive |
| Spleen | Atlantic salmon | 1 | PRJNA72713 | BioProject |
| Skin | Atlantic salmon | 1 | PRJNA72713 | BioProject |

**Supplementary Table 2.** Phylogeny-based annotation of glycoside hydrolase family 18- and chitin synthase genes and proteins in Atlantic salmon.

| **Clade** | **Gene name used in this paper** | **NCBI gene ID** | **Protein name used in this paper** | **RefSeq protein accession number** |
| --- | --- | --- | --- | --- |
| CHIA | *chia.3* | 106565088 | Chia.3 | XP_013987246.1 |
|  | *chia.4* | 106565087 | Chia.4 | XP_013987245.1 |
|  | *chia.7* | 106572401 | Chia.7 | XP_014002002.1 |
| CHIO I | *chia.2* | 106565093 | Chia.2 | XP_013987254.1 |
|  | *chia.9* | 106583514 | Chia.9 | XP_014023261.1 |
|  | *chia.8* | 106577510 | Chia.8 | XP_014011074.1 |
|  | *chia.6* | 106567267 | Chia.6 | XP_013991841.1 |
| CHIO II | *chia.1* | 106565309 | Chia.1 | XP_013987730.1 |
|  | *chia.10* | 106583513 | Chia.10 | XP_014023260.1 |
| CHIT1 | *chia.5* | 106567333 | Chia.5 | XP_013991941.1 |
| CHID1 | *chid1* | 100194776 | Chid1 | NP_001133277.1 |
| CTBS | *ctbs* | 100195549 | Ctbs | XP_014071095.1 |
| CHS | *chs1a* | 106589822 | Chs1a | XP_014035674.1 |
|  | *chs1b* | 106612488 | Chs1b | XP_014069148.1 |
|  | *chs2* | 106562657 | Chs2 | XP_013983103.1 |
|  | *chs3* | 106570137 | Chs3 | XP_013997648.1 |

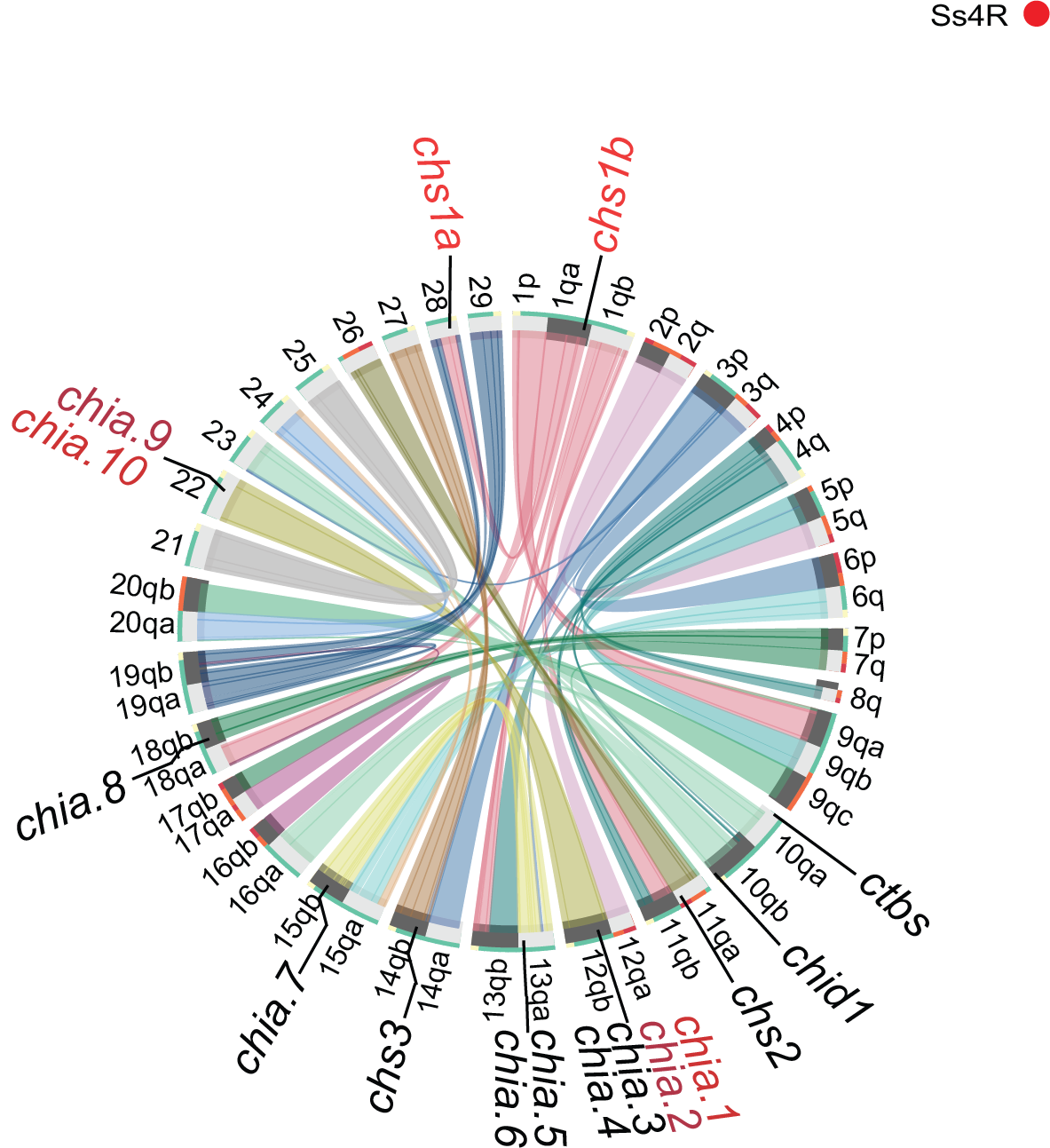

**Supplementary Figure 1.** Circos plot showing the chromosomal location of glycoside hydrolase family 18 and chitin synthase genes in Atlantic salmon. The blocks represent homologous regions in the genome with the 29 Atlantic salmon chromosomes, see [(Lien *et al.* 2016)](https://sciwheel.com/work/citation?ids=1436568&pre=&suf=&sa=0&dbf=0) for more information. The genes marked in red (*chia.1 + chia.10, chia.2 + chia.9, chs1a + chs1b*) are located on homologous regions of different chromosomes and are hypothesized to result from the salmonid-specific whole-genome duplication event (Ss4R).

**Supplementary Table 3.** Results from the KEGG enrichment with pathways, number of genes, p-value, NCBI gene ID, and gene description.

| **KEGG Pathway** | **Genes** | **Genes in list** | **P-value** | **PathwayID** | **NCBI Gene ID** | **NCBI Gene description** |
| --- | --- | --- | --- | --- | --- | --- |
| Amino sugar and nucleotide sugar metabolism | 84 | 6 | 8.79E-10 | path:sasa00520 | 106560290 | UDP-N-acetylhexosamine pyrophosphorylase-like |
| Amino sugar and nucleotide sugar metabolism | 84 | 6 | 8.79E-10 | path:sasa00520 | 106565093 | Acidic mammalian chitinase-like (*chia.2*) |
| Amino sugar and nucleotide sugar metabolism | 84 | 6 | 8.79E-10 | path:sasa00520 | 106565309 | Acidic mammalian chitinase-like (*chia.1*) |
| Amino sugar and nucleotide sugar metabolism | 84 | 6 | 8.79E-10 | path:sasa00520 | 106567267 | Acidic mammalian chitinase-like (*chia.6*) |
| Amino sugar and nucleotide sugar metabolism | 84 | 6 | 8.79E-10 | path:sasa00520 | 106577510 | Acidic mammalian chitinase-like (*chia.8*) |
| Amino sugar and nucleotide sugar metabolism | 84 | 6 | 8.79E-10 | path:sasa00520 | 106612488 | Uncharacterized (*chs1b*) |
| Metabolic pathways | 2355 | 13 | 2.43E-06 | path:sasa01100 | 101448035 | beta-carotene 15,15'-monooxygenase 1 like |
| Metabolic pathways | 2355 | 13 | 2.43E-06 | path:sasa01100 | 106560290 | UDP-N-acetylhexosamine pyrophosphorylase-like |
| Metabolic pathways | 2355 | 13 | 2.43E-06 | path:sasa01100 | 106564978 | beta-1,3-galactosyl-O-glycosyl-glycoprotein beta-1,6-N-acetylglucosaminyltransferase 3-like |
| Metabolic pathways | 2355 | 13 | 2.43E-06 | path:sasa01100 | 106565093 | Acidic mammalian chitinase-like (*chia.2*) |
| Metabolic pathways | 2355 | 13 | 2.43E-06 | path:sasa01100 | 106565309 | Acidic mammalian chitinase-like (*chia.1*) |
| Metabolic pathways | 2355 | 13 | 2.43E-06 | path:sasa01100 | 106567267 | Acidic mammalian chitinase-like (*chia.6*) |
| Metabolic pathways | 2355 | 13 | 2.43E-06 | path:sasa01100 | 106572530 | diacylglycerol kinase alpha-like |
| Metabolic pathways | 2355 | 13 | 2.43E-06 | path:sasa01100 | 106575817 | acyl-CoA dehydrogenase, long chain |
| Metabolic pathways | 2355 | 13 | 2.43E-06 | path:sasa01100 | 106576053 | aldose reductase-like |
| Metabolic pathways | 2355 | 13 | 2.43E-06 | path:sasa01100 | 106577510 | Acidic mammalian chitinase-like (*chia.8*) |
| Metabolic pathways | 2355 | 13 | 2.43E-06 | path:sasa01100 | 106609639 | ST8 alpha-N-acetyl-neuraminide alpha-2,8-sialyltransferase 1 |
| Metabolic pathways | 2355 | 13 | 2.43E-06 | path:sasa01100 | 106609667 | alpha-N-acetylneuraminide alpha-2,8-sialyltransferase-like |
| Metabolic pathways | 2355 | 13 | 2.43E-06 | path:sasa01100 | 106612488 | Uncharacterized (*chs1b*) |
| Glycosphingolipid biosynthesis - ganglio series | 29 | 2 | 0.00059208 | path:sasa00604 | 106609639 | ST8 alpha-N-acetyl-neuraminide alpha-2,8-sialyltransferase 1 |
| Glycosphingolipid biosynthesis - ganglio series | 29 | 2 | 0.00059208 | path:sasa00604 | 106609667 | alpha-N-acetylneuraminide alpha-2,8-sialyltransferase-like |
| Glycosphingolipid biosynthesis - globo and isoglobo series | 30 | 2 | 0.00063388 | path:sasa00603 | 106609639 | ST8 alpha-N-acetyl-neuraminide alpha-2,8-sialyltransferase 1 |
| Glycosphingolipid biosynthesis - globo and isoglobo series | 30 | 2 | 0.00063388 | path:sasa00603 | 106609667 | alpha-N-acetylneuraminide alpha-2,8-sialyltransferase-like |
| Glycosphingolipid biosynthesis - lacto and neolacto series | 46 | 2 | 0.00148954 | path:sasa00601 | 106609639 | ST8 alpha-N-acetyl-neuraminide alpha-2,8-sialyltransferase 1 |
| Glycosphingolipid biosynthesis - lacto and neolacto series | 46 | 2 | 0.00148954 | path:sasa00601 | 106609667 | alpha-N-acetylneuraminide alpha-2,8-sialyltransferase-like |
| Retinol metabolism | 74 | 2 | 0.00380361 | path:sasa00830 | 100196171 | putative all-trans-retinol 13,14-reductase |
| Retinol metabolism | 74 | 2 | 0.00380361 | path:sasa00830 | 101448035 | beta-carotene 15,15'-monooxygenase 1 like |
| Glycerolipid metabolism | 100 | 2 | 0.00683169 | path:sasa00561 | 106572530 | diacylglycerol kinase alpha-like |
| Glycerolipid metabolism | 100 | 2 | 0.00683169 | path:sasa00561 | 106576053 | aldose reductase-like |

[**References**](https://sciwheel.com/work/bibliography?atCursor=false)

[Lien, S., B. F. Koop, S. R. Sandve, J. R. Miller, M. P. Kent *et al.*, 2016 The Atlantic salmon genome provides insights into rediploidization. Nature 533: 200–205.](https://sciwheel.com/work/bibliography/1436568)
